# Segmentation pathway genes in the Asian citrus psyllid, *Diaphorina citri*

**DOI:** 10.1101/2020.12.24.424320

**Authors:** Sherry Miller, Teresa D. Shippy, Prashant S Hosmani, Mirella Flores-Gonzalez, Lukas A Mueller, Wayne B Hunter, Susan J Brown, Tom D’elia, Surya Saha

## Abstract

Insects have a segmented body plan that is established during embryogenesis when the anterior-posterior (A-P) axis is divided into repeated units by a cascade of gene expression. The cascade is initiated by protein gradients created by translation of maternally provided mRNAs, localized at the anterior and posterior poles of the embryo. Particular combinations of these proteins activate specific gap genes to divide the embryo into distinct regions along the A-P axis. Gap genes then activate pair-rule genes, which are usually expressed in part of every other segment. The pair-rule genes, in turn, activate expression of segment polarity genes in a portion of each segment. The segmentation genes are generally conserved among insects, although there is considerable variation in how they are deployed. We annotated 24 segmentation gene homologs in the Asian citrus psyllid, *Diaphorina citri*. We identified most of the genes that were expected to be present based on known phylogenetic distribution. Two exceptions were *eagle* and *invected*, which are present in at least some hemipterans, but were not identified in *D. citri*. Many of these genes are likely to be essential for *D. citri* development and thus may be useful targets for pest control methods.

## Introduction

Segmentation is the process by which repeated units of similar groups of cells are created along the anterior-posterior axis of a developing embryo. The molecular mechanisms involved in this process were first elucidated by large scale developmental mutant screens in the insect model *Drosophila melanogaster* [1–5]. In *Drosophila*, segmentation begins with cytoplasmic inheritance of mRNAs that are maternally produced and provided to the oocyte. The products of these maternal-effect genes create gradients that define positional information within the embryo and activate a group of genes known as gap genes. Gap genes are expressed in broad, well-defined domains in the early embryo and activate the next set of transcription factors, the pair-rule genes. Pair-rule genes are expressed in every other segment of the developing embryo and together they activate the expression of segment polarity genes which are expressed in every segment of the developing embryo. Comparative studies in diverse arthropod species have shown that some aspects of the segmentation pathway are highly conserved while other aspects have undergone evolutionary change [6]. The hemipteran insects that have been examined seem to employ a particularly divergent method of segmentation. Most strikingly, the pair-rule genes, generally considered the most conserved portion of the segmentation pathway among insects, have lost their pair-rule expression and function and are expressed segmentally in at least some hemipterans [7–9].

We are participating in a community annotation project to annotate the genome of the hemipteran agricultural pest, *Diaphorina citri*, also known as the Asian citrus psyllid. *D. citri* is the vector responsible for the spread of Huanglongbing (citrus greening disease) which has devastated the citrus industry. Here we describe the identification and annotation of *D. citri* orthologs of genes identified in *Drosophila* segmentation. We found and manually annotated 24 homologs of these genes, as well as several related genes. In most cases, the presence or absence of particular genes in *D. citri* is consistent with expectations based on reports in other insects. However, eagle and invected, two segmentation genes expected to be present, appear to be missing from the *D. citri* v3 genome. Our annotations pave the way for future work aimed at understanding the expression and function of these genes during *D. citri* segmentation and the identification of essential genes that could be used as insect control targets.

## Results and Discussion

We searched the *D. citri* v3 genome for orthologs of genes known to be involved in segmentation in *Drosophila* [10] (Table 1). We then used available evidence to manually annotate the genes that were present [11,12] (Table 1–2). Most manual annotations were straightforward and performed using our workflow [see Methods], so only those requiring additional explanation are described in detail here.

**Table 1.**
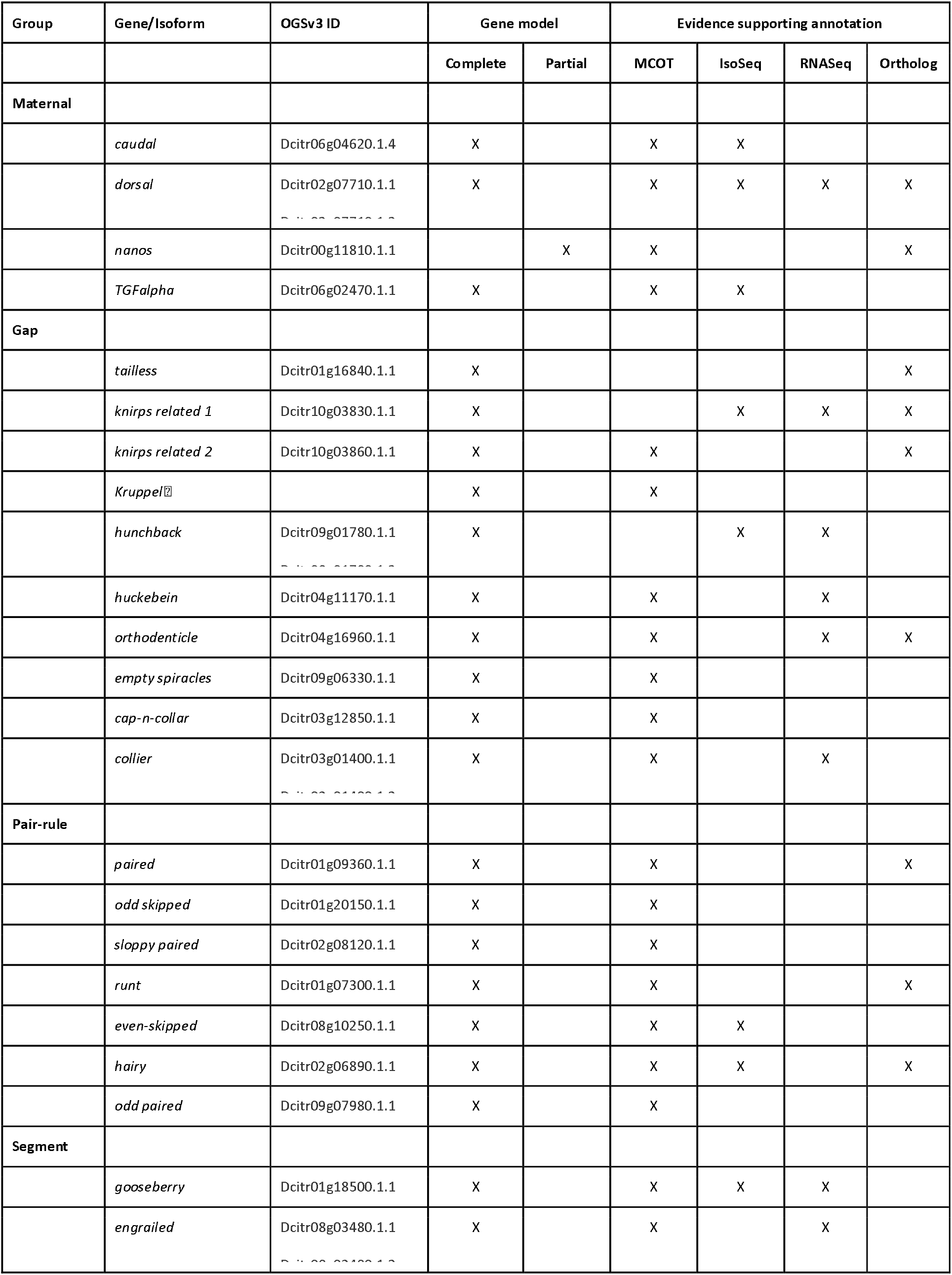
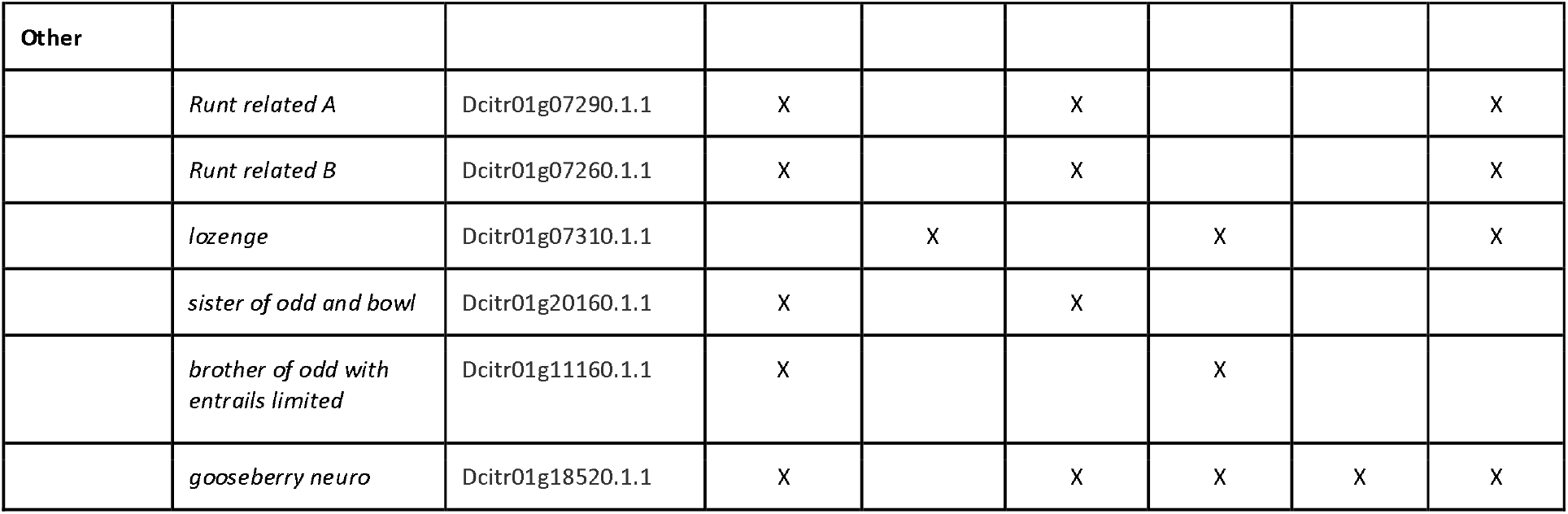
Annotated *D. citri* segmentation gene orthologs. Each manually annotated gene has been assigned an OGS3 gene identifier and denoted as a partial or complete model. Evidence used for manual annotation was also recorded. MCOT [13] and IsoSeq [11] indicate use of these genome-independent transcriptomes to validate the annotation. RNASeq indicates use of mapped RNASeq reads, while orthologs indicates comparison to related proteins was essential for the annotation. Additional information on specific evidence types is included in [12]. ⍰denotes genes that were complete in OGS2 but are partial in OGS3.

Annotated *D. citri* segmentation gene orthologs
| Group | Gene/Isoform | OGSv3 ID | Gene model |  | Evidence supporting annotation |  |  |  |
| --- | --- | --- | --- | --- | --- | --- | --- | --- |
|  |  |  | Complete | Partial | MCOT | IsoSeq | RNASeq | Ortholog |
| Maternal |  |  |  |  |  |  |  |  |
|  | *caudal* | Dcitr06g04620.1.4 | X |  | X | X |  |  |
|  | *dorsal* | Dcitr02g07710.1.1 | X |  | X | X | X | X |
|  | *nanos* | Dcitr00g11810.1.1 |  | X | X |  |  | X |
|  | *TGFalpha* | Dcitr06g02470.1.1 | X |  | X | X |  |  |
| Gap |  |  |  |  |  |  |  |  |
|  | *tailless* | Dcitr01g16840.1.1 | X |  |  |  |  | X |
|  | *knirps related 1* | Dcitr10g03830.1.1 | X |  |  | X | X | X |
|  | *knirps related 2* | Dcitr10g03860.1.1 | X |  | X |  |  | X |
|  | *Kruppel* |  | X |  | X |  |  |  |
|  | *hunchback* | Dcitr09g01780.1.1 | X |  |  | X | X |  |
|  | *huckebein* | Dcitr04g11170.1.1 | X |  | X |  | X |  |
|  | *orthodenticle* | Dcitr04g16960.1.1 | X |  | X |  | X | X |
|  | *empty spiracles* | Dcitr09g06330.1.1 | X |  | X |  |  |  |
|  | *cap-n-collar* | Dcitr03g12850.1.1 | X |  | X |  |  |  |
|  | *collier* | Dcitr03g01400.1.1 | X |  | X |  | X |  |
| Pair-rule |  |  |  |  |  |  |  |  |
|  | *paired* | Dcitr01g09360.1.1 | X |  | X |  |  | X |
|  | *odd skipped* | Dcitr01g20150.1.1 | X |  | X |  |  |  |
|  | *sloppy paired* | Dcitr02g08120.1.1 | X |  | X |  |  |  |
|  | *runt* | Dcitr01g07300.1.1 | X |  | X |  |  | X |
|  | *even-skipped* | Dcitr08g10250.1.1 | X |  | X | X |  |  |
|  | *hairy* | Dcitr02g06890.1.1 | X |  | X | X |  | X |
|  | *odd paired* | Dcitr09g07980.1.1 | X |  | X |  |  |  |
| Segment |  |  |  |  |  |  |  |  |
|  | *gooseberry* | Dcitr01g18500.1.1 | X |  | X | X | X |  |
|  | *engrailed* | Dcitr08g03480.1.1 | X |  | X |  | X |  |

Table 1. Each manually annotated gene has been assigned an OGS3 gene identifier and denoted as a partial or complete model.
| Other |  |  |  |  |  |  |  |  |
| --- | --- | --- | --- | --- | --- | --- | --- | --- |
|  | *Runt related A* | Dcitr01g07290.1.1 | X |  | X |  |  | X |
|  | *Runt related B* | Dcitr01g07260.1.1 | X |  | X |  |  | X |
|  | *lozenge* | Dcitr01g07310.1.1 |  | X |  | X |  | X |
|  | *sister of odd and bowl* | Dcitr01g20160.1.1 | X |  | X |  |  |  |
|  | *brother of odd with entrails limited* | Dcitr01g11160.1.1 | X |  |  | X |  |  |
|  | *gooseberry neuro* | Dcitr01g18520.1.1 | X |  | X | X | X | X |

**Table 2.**
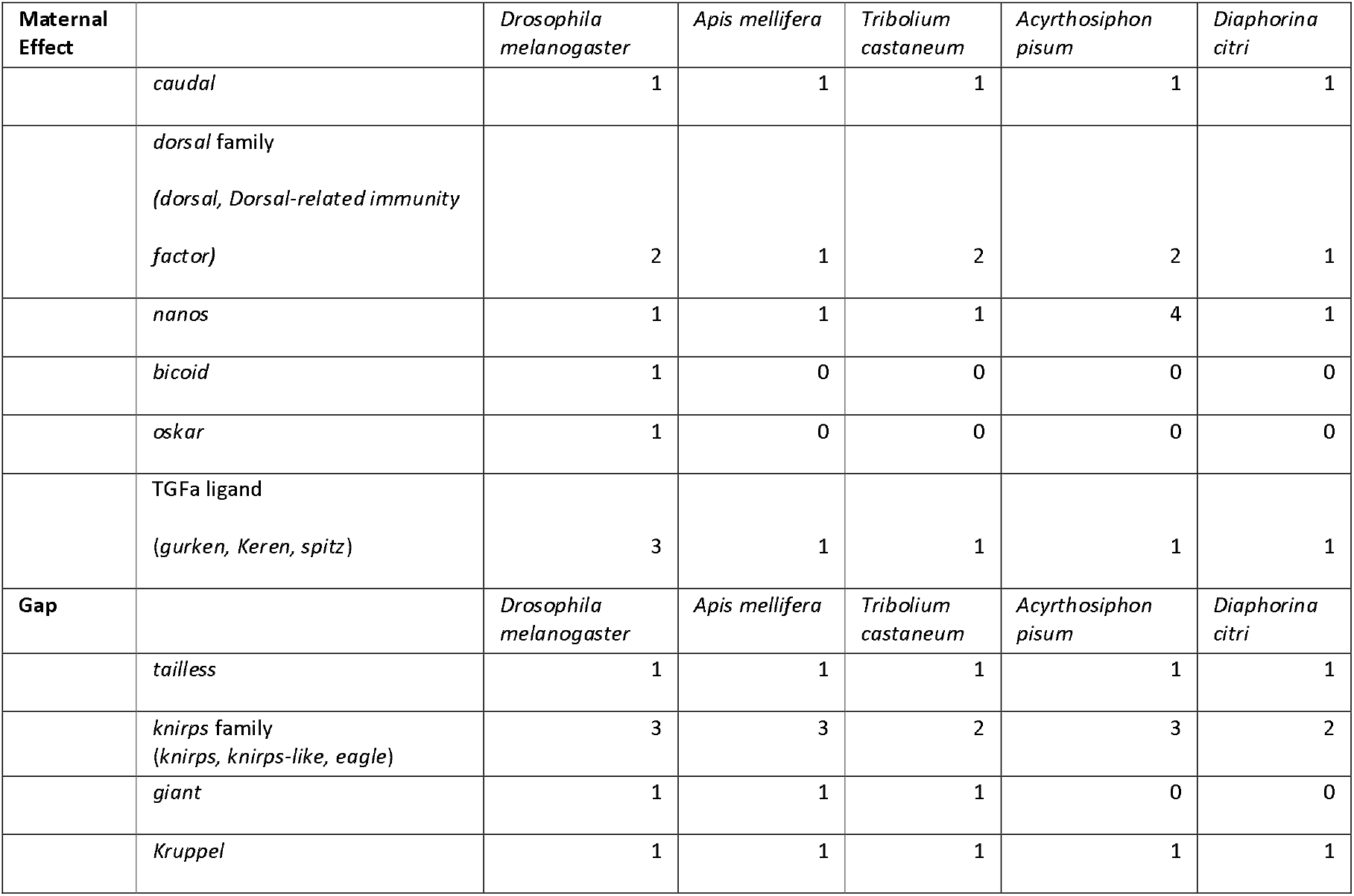
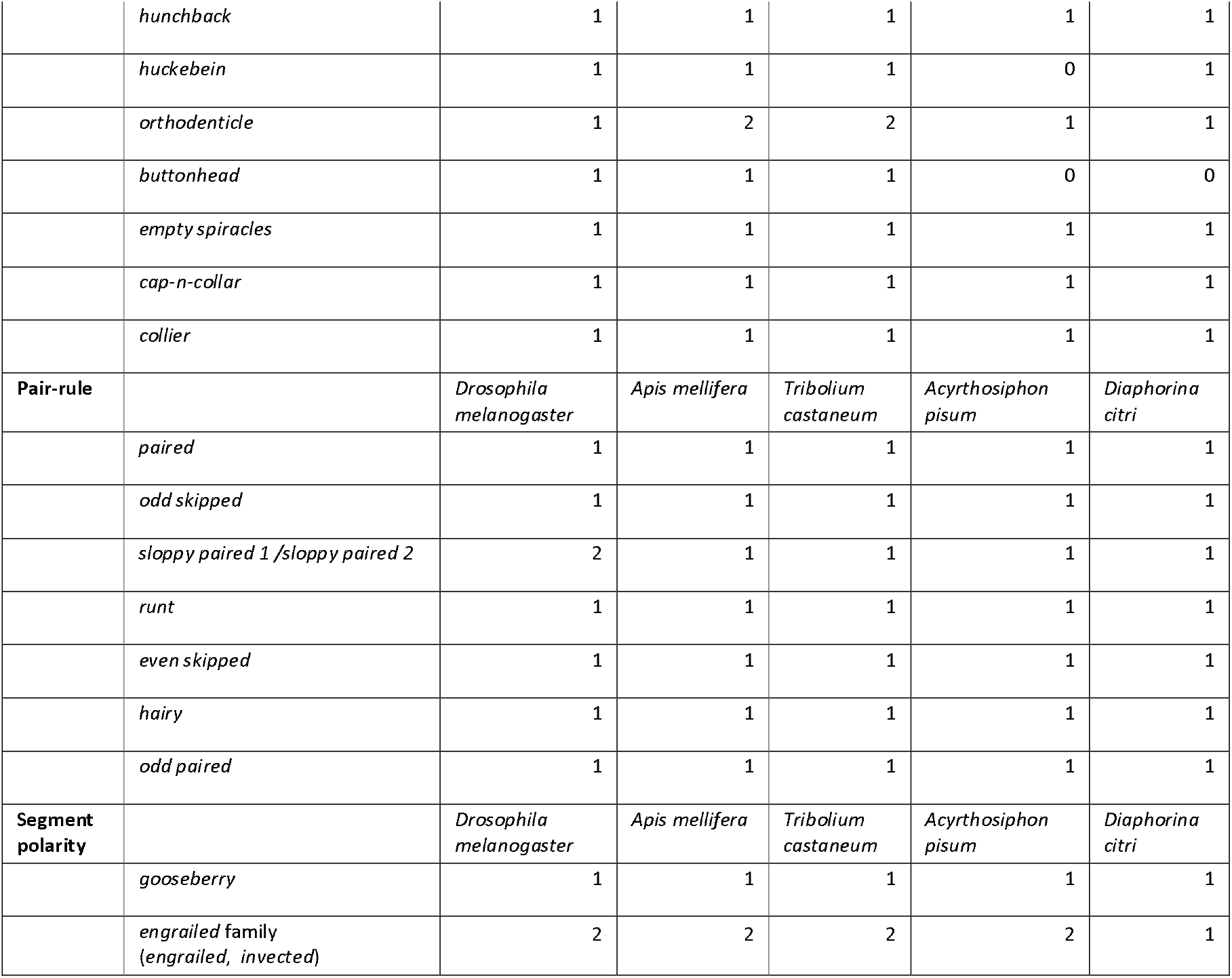
Segmentation gene ortholog number. The *Drosophila melanogaster* numbers were determined from Flybase. Ortholog numbers for *Apis mellifera* [18], *Tribolium castaneum* [40] and *Acyrthosiphon pisum* [17] are based on genome publications or NCBI. *Diaphorina citri* ortholog numbers represent our final manual annotation.

### Maternal effect genes

One-to-one orthologs of *caudal* (*cad*), *dorsal* (*dl*), and *nanos* (*nos*) were found in the *D. citri* v3 genome. *dl* was previously annotated in the *D. citri* genome v1.1 because of its role in innate immunity [13]. Here we annotated a second isoform of dl (Table 1).

We also identified a single TGF-α ligand-encoding gene in *D. citri*. In *Drosophila*, there are three *TGF-α* ligand paralogs (*grk, Krn, spi*), but in many other insects, only one *TGF-α* ligand has been identified [14–18]. Orthologs for *bicoid* (*bcd*) and *oskar* (*osk*) were not found in *D. citri* (Table 2), which is consistent with the previously described phylogenetic distribution of these genes [19].

### Gap genes

One-to-one orthologs of the gap genes *tailless* (*tll*), *Kruppel* (*Kr*), *hunchback* (*hb*) *huckebein* (*hkb*), *empty spiracles* (*ems*), *cap-n-collar* (*cnc*), and *collier* (*col*) were identified and annotated in the *D. citri* v3 genome (Tables 1–2). Except for *hkb*, which was reported missing in *A. pisum* [17], conservation of these genes was expected (Table 2). Two *knirps-related* (*knrl*) genes and two *orthodenticle* (*otd*) homologs were annotated and are discussed in more detail below. *giant* (*gt*) and *buttonhead* (*btd*) appear to be absent in the v3 *D. citri* genome assembly (Table 2).

Phylogenetics of the *knirps* family suggests that a single ancestral gene duplicated early in the insect lineage producing two paralogs that have been called *knirps-related* (*knrl*) and eagle (eg) [20]. Subsequent duplications have occurred in various insect lineages. A duplication in the lineage leading to Drosophila resulted in the paralogs *knirps* and *knirps-like* (also called *knirps-related*) [20]. A separate duplication of *knrl* seems to have occurred in the hemipteroid lineage leading to three *knirps* family genes (two *knrl* and one *eg*) in most hemipterans [20]. However, in the *D. citri* genome v3 we were only able to identify two knirps family genes (Tables 1–2), one of which was annotated as *knirps* in *D. citri* genome v1.1 [13]. The two *knirps* family genes are located on the same chromosome, about 400 kb apart. Both predicted proteins contain the highly conserved 94 amino acid N terminal domain and the C terminal PIDLS motif commonly found in *knirps* family members [20]. However, neither contains the GASS-domain motif that is unique to the Eg protein [20]. Due to the lack of this signature Eagle motif, the resulting *D. citri* annotations were named *knirps related 1* (*knrl1*) and *knirps related 2* (*knrl2*). Despite the lack of the GASS-domain, it is possible that *D. citri knrl2* is the ortholog of eg since phylogenetic analysis was inconclusive (data not shown). Interestingly, *D. citri knrl1* has a small exon just 5’ of the highly conserved coding exon that is the first exon in most *knirps* family genes (Figure 1). Similar gene structure has been reported for one *knirps* family gene each in *D. melanogaster*, the honeybee *Apis mellifera, A. pisum*, and the human louse *Pediculus* [20]. All but *D. melanogaster* share small stretches of sequence identity in the amino acid sequence encoded by this additional exon, suggesting that the 5’ exon might have been present in a common ancestor. This model suggests a duplication and acquisition of an additional exon by one paralog early in the insect lineage. The paralog containing the additional exon appears to have been lost sometime after the divergence of the Hymenoptera from the rest of the Holometabola.

**Figure 1.**
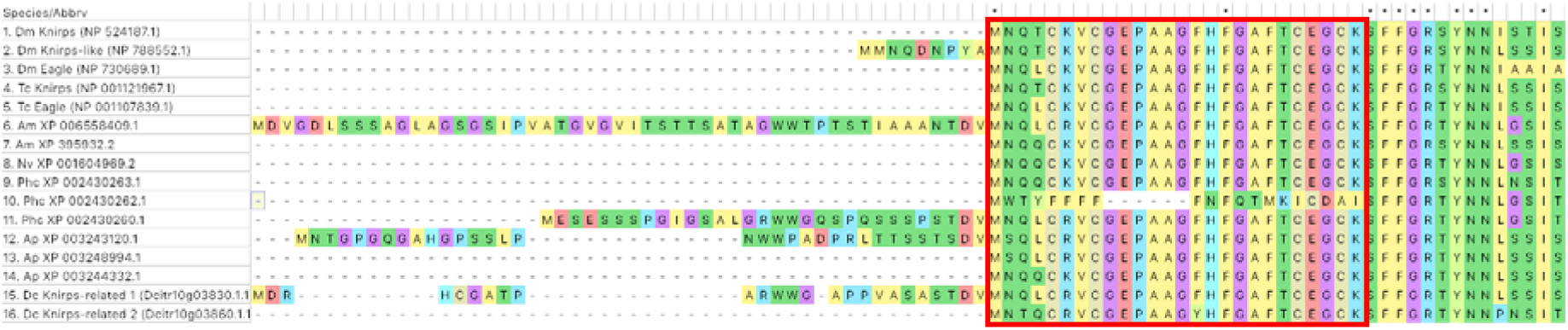
Multiple alignment of the N-terminal region of Knirps family proteins. The red box denotes the sequence encoded by a highly conserved exon [20]. Species represented are *Drosophila melanogaster* (Dm), *Tribolium castaneum* (Tc), *Apis mellifera* (Am), *Nasonia vitripennis* (Nv), *Pediculus humanus* (Ph), *Acyrthosiphon pisum* (Ap) and *Diaphorina citri* (Dc). Named proteins have been manually annotated, while those with only an accession number are computationally predicted. Five of the *knirps* family genes (one each in Dm, Am, Ph, Ap and Dc) have sequence upstream of the universally conserved sequence that typically begins in the first exon [20 and this work]. In these proteins the conserved core sequence (red box) begins in the second exon. Four of the five proteins with an additional 5’ exon (Am, Ph, Ap and Dc) share a small region of sequence identity at the C-terminal end of the sequence encoded by this exon.

Most insects have two *otd* genes. However, in both *Drosophila* and *A. pisum* only one *otd* gene has been identified. *Drosophila* is missing the *otd-2* ortholog, while *otd-1* has apparently been lost in pea aphids [21]. In the *D. citri* v3 genome, we found two *otd* genes adjacent to one another on chromosome 4 (Table 1). Phylogenetic analysis suggests that one of these genes is an *otd-1* ortholog, while the other is an *otd-2* ortholog (Figure 2). *otd-1* and *otd-2* are also clustered in other insects and crustaceans where their genomic location has been examined [22], suggesting that there are probably evolutionary constraints preventing their separation.

**Figure 2.**
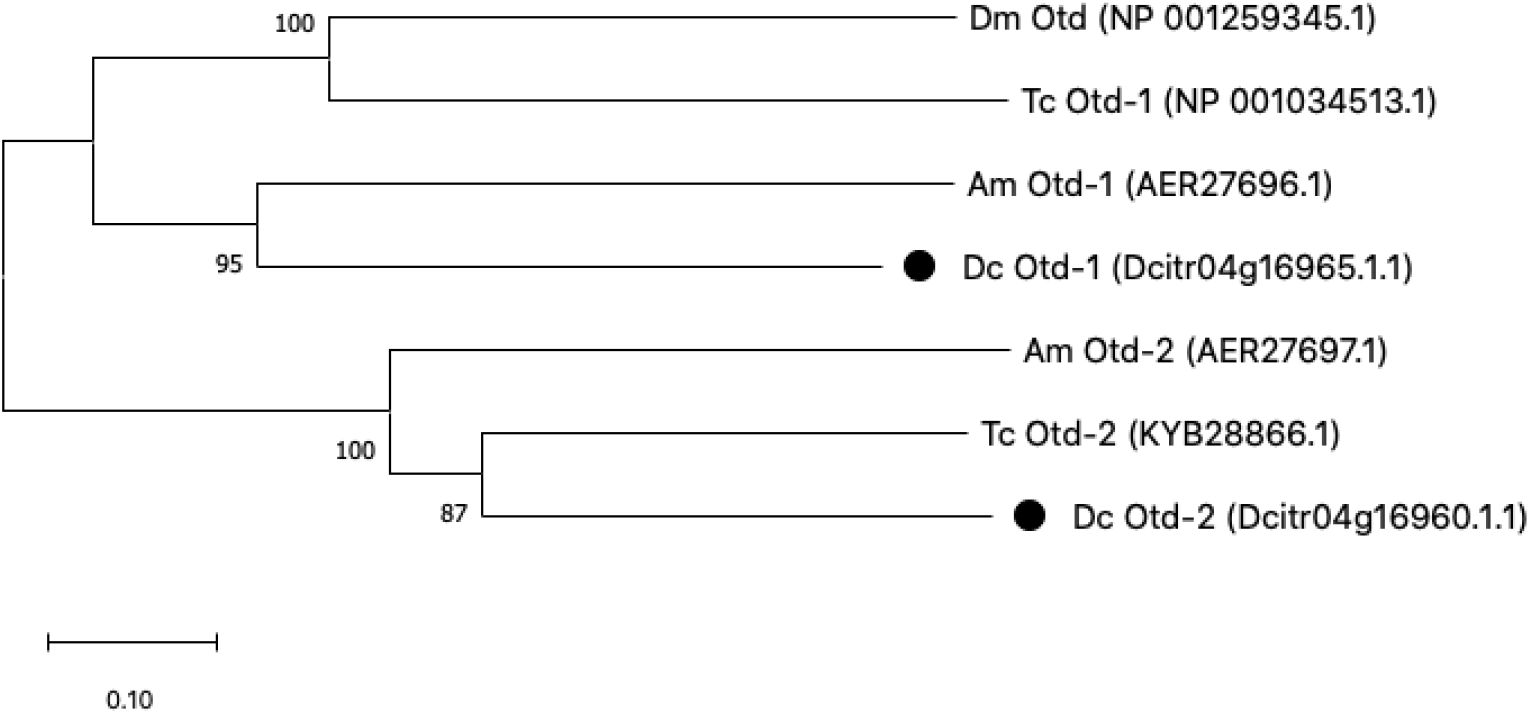
Neighbor joining tree of Otd homologs from Drosophila melanogaster (Dm), Tribolium castaneum (Tc), Apis mellifera (Am), and Diaphorina citri (Dc). *D. citri* proteins are marked with black circles.

*gt* is conserved in some, but not all, hemipterans. A gt ortholog was not found in the *A. pisum* genome [17], however, apparent gt orthologs are present in *Rhodnius prolixus, Cimex lectularius, Halyomorpha halys, Oncopeltus fasciatus* and *Bemisia tabaci* [23–27]. RNAi studies indicate that the *R. prolixus* and *O. fasciatus* gt orthologs both function as gap genes [23,26]. We performed BLAST searches of *D. citri* genome v3 with all of these hemipteran Gt orthologs, but were unable to identify a *D. citri* gt ortholog (Table 2).

*btd* is a member of the Sp-family of transcription factors. Recent reports indicate that presence of three Sp members is likely the ancestral state for arthropods and perhaps all metazoans [28]. These three Sp-family genes cluster into three monophyletic clades (Sp5/btd, Sp1-4/ (Sp-pps) and Sp6-9 (Sp1)) [28]. Despite the fact that three Sp-family members appears to be the ancestral state, *btd* is absent from the *A. pisum* genome [17] and repeated efforts to clone *btd* from *Oncopeltus fasciatus* has only resulted in the identification of the two non *btd* Sp genes, suggesting that *btd* may have been lost in the lineage leading to hemipterans. We too were unable to find a true *btd* ortholog in either the *D. citri* genome v3 or in independent *de novo* transcriptomes (Table 2), but we did find two Sp-family members that appear to be orthologous to *Sp1* and *Spps*.

### Pair-rule genes

One-to-one orthologs were found for all pair-rule genes examined, including *paired* (*prd*), *odd skipped* (*odd*), *sloppy paired* (*slp*), *runt* (*run*), *even skipped* (*eve*), *hairy* (*h*) and *odd paired* (*opa*) in the *D. citri* genome v3 (Tables 1–2). A partial copy of h had been annotated in a previous genome version [13]. Three of the pair-rule genes we annotated have closely related paralogs and required additional analysis before gene identities could be assigned. Prd is a member of the Pax3/7 family of proteins. In *Drosophila* there are three Pax3/7 family genes which are known to play a role in segmentation and neurogenesis, *prd, gooseberry* (*gsb*) and *gooseberry-neuro* (*gsb-n*). While the number of Pax3/7 genes varies in arthropods, data from insects and arachnids suggest that the Pax3/7 roles in segmentation and neurogenesis are likely to be conserved in all arthropods [29]. In the *D. citri* genome v3 we also found three Pax3/7 genes that we named *D.citri_paired*, *D. citri_gooseberry* and *D. citri_gooseberry-neuro* based on reciprocal BLAST analysis and genomic location (Tables 1–2). The *gsb* and *gsb-n* orthologs are discussed in more detail in the segment polarity gene section.

*odd* is a zinc finger transcription factor that has three close relatives known as *brother of odd with entrails limited* (*bowl*), *sister of odd and bowl* (*sob*) and *drumstick* (*drum*). All four genes are located in an evolutionarily conserved cluster [30]. In *D. citri* genome v3, *drum*, *odd* and *sob* are all located within 400 kb, with *odd* and *sob* overlapping one another on opposite strands. It’s not clear whether the overlap is correct or results from misassembly, but the genes are almost certainly located very close together. *D.citri bowl* is located on the same chromosome about 20 Mb away. Separation of *bowl* from the rest of the cluster has also been observed in *Anopheles gambiae* [31].

There are four *runt* domain-containing genes in insects: *run, Runt related A* (*RunxA*), *Runt related B* (*RunxB*) and *lozenge* (*lz*). All four genes are typically found in a cluster and their order and orientation is well conserved across insects [18] (Figure 3). We were able to annotate full length models for all four genes in the *D. citri* genome. It appears that the cluster is intact, with all four RDP genes identified in their expected order within a 300 kb region (Figure 3).

**Figure 3.**
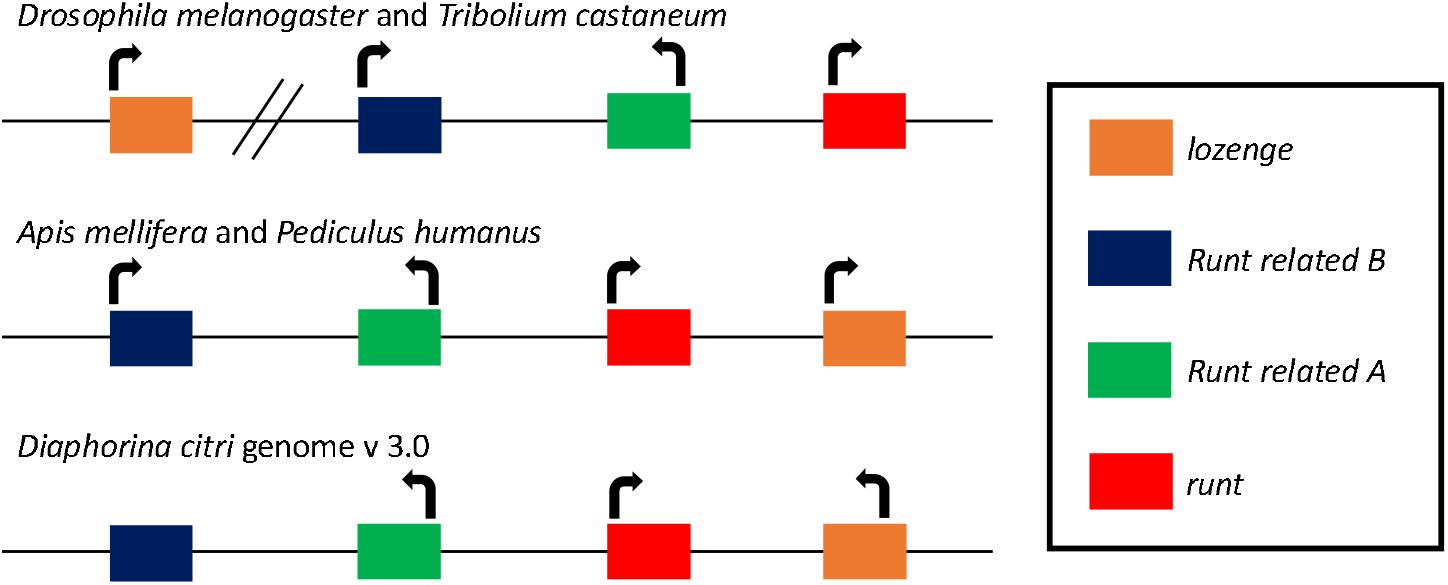
Runt Domain (RD) cluster in representative insects. Cluster information from other insects was obtained from [48]. The RD clusters in D. melanogaster and T. castaneum have three genes in a core cluster, with lozenge (lz) separated from the cluster, but on the same chromosome distal to runt (run). The RD clusters in A. mellifera and P. humanus have all four genes clustered together with lz proximal to run. The RD cluster in D. citri most closely follows the pattern seen in A. mellifera and P. humanus. D. citri lz appears to be transcribed in the opposite direction compared to other insects, but it is possible that this is due to local misassembly. The orientation of D. citri Runt related B (RunxB) is uncertain, since there are tandem artifactual duplicates that are on opposite strands. We chose the RunxB copy closest to Runt related A (RunxA) for annotation. Future assembly improvements may help resolve the gene orientation in this cluster.

### Segment polarity genes

Many segment polarity genes are members of the Wnt and Hedgehog signaling pathways. The manual annotation of the Wnt pathway genes in the *D. citri* genome v3 is described in a separate report [32]. Here we report the manual annotation of the segment polarity genes *gooseberry* (*gsb*) and *engrailed* (*en*) (Table 1). *gsb* and *en* each have a tightly linked paralog in many insects [33–35]. Surprisingly, we were unable to find the *en* paralog invected (*inv*) in the current genome assembly or the *de novo* transcriptome. We did find and annotate the *gsb* paralog *gooseberry-neuro* (*gsb-n*) in its expected position adjacent to *gsb* (Table 1). This positional information helped verify the identity of *gsb-n*, since phylogenetic analysis was inconclusive.

## Conclusion

We searched for orthologs of 33 Drosophila segmentation genes in the *D. citri* v3 genome and identified and annotated 24 homologous genes. We were unable to find orthologs for 10 of the *Drosophila* genes, while *D. citri* has one segmentation gene (*otd-2*) whose ortholog has been lost in *Drosophila*. Most of these absences, except *eagle* and *invected*, were expected based on the known phylogenetic distribution of the genes. While all the genes discussed in this report were initially identified because of their role in embryonic patterning and segmentation in *Drosophila*, many of them also have other important functions such as pole cell development, neural stem cell maintenance, sex determination, and immune function. Analysis of expression patterns and gene function will be required to determine which of these genes are involved in *D. citri* segmentation, and which might be good targets for control of *D. citri* and Huanglongbing.

## Methods

Segmentation gene orthologs in *D. citri* genome v3 [11] were identified by BLAST search and confirmed by reciprocal BLAST. Manual annotation was performed in Apollo 2.1.0 [36] using available evidence, such as RNA-Seq reads, IsoSeq transcripts and *de novo*-assembled transcripts. Multiple alignments were performed with MUSCLE [37] or MEGA7 [38] and phylogenetic analysis was done in MEGA7 or MEGA X [39]. A list of orthologs used in these analyses can be found in Table 3. More details of the annotation process are described in [12].

**Table 3.**
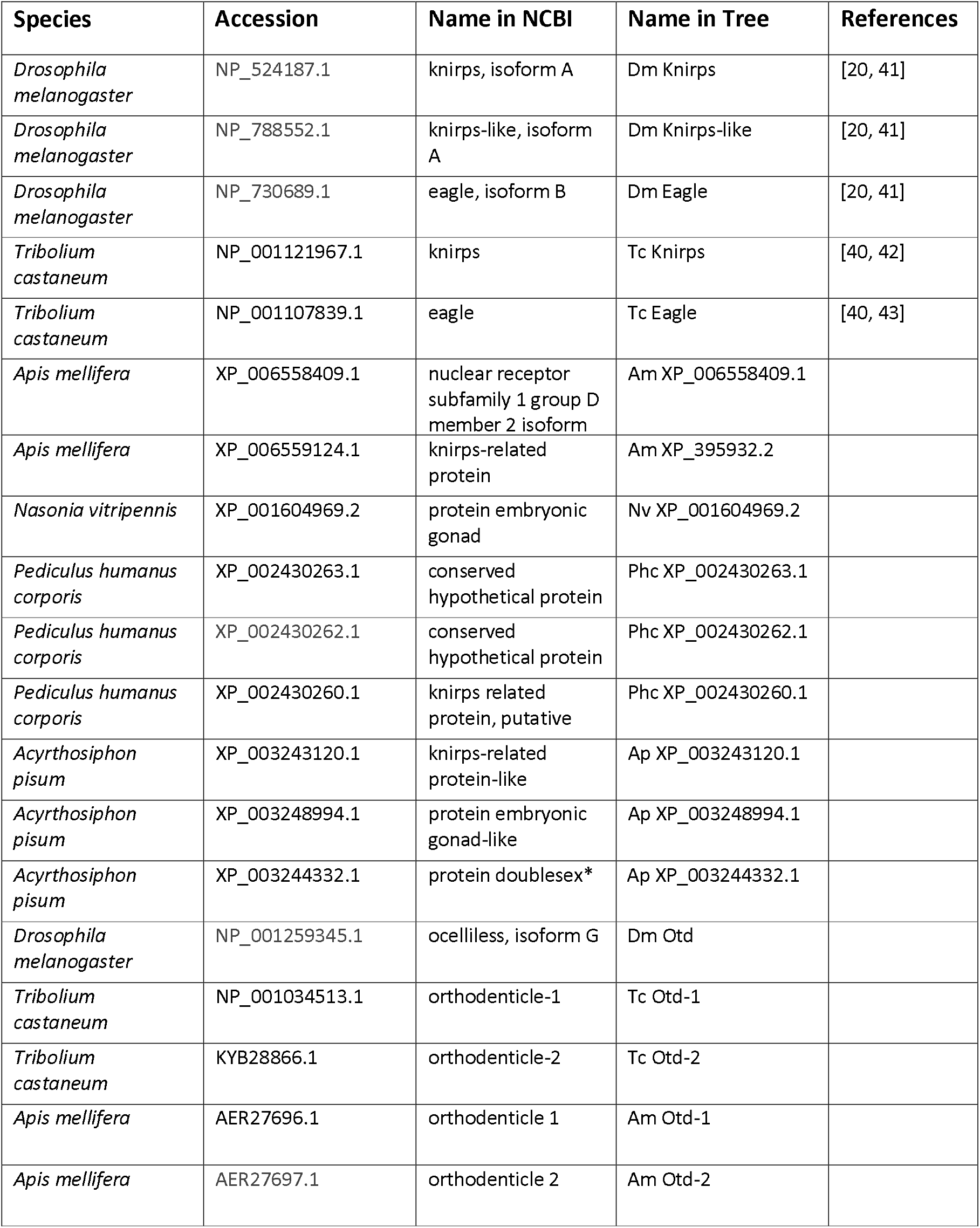
Orthologs used in Alignments and Phylogenetic Analysis Species, accession number, full name and abbreviated name are provided for all orthologs used multiple alignments and phylogenetic trees [20,40–47]. Gene names from NCBI that are marked with an asterisk (*) appear to be incorrect annotations.

| Species | Accession | Name in NCBI | Name in Tree | References |
| --- | --- | --- | --- | --- |
| *Drosophila melanogaster* | NP_524187.1 | knirps, isoform A | Dm Knirps | [20, 41] |
| *Drosophila melanogaster* | NP_788552.1 | knirps-like, isoform A | Dm Knirps-like | [20, 41] |
| *Drosophila melanogaster* | NP_730689.1 | eagle, isoform B | Dm Eagle | [20, 41] |
| *Tribolium castaneum* | NP_001121967.1 | knirps | Tc Knirps | [40, 42] |
| *Tribolium castaneum* | NP_001107839.1 | eagle | Tc Eagle | [40, 43] |
| *Apis mellifera* | XP_006558409.1 | nuclear receptor subfamily 1 group D member 2 isoform | Am XP_006558409.1 |  |
| *Apis mellifera* | XP_006559124.1 | knirps-related protein | Am XP_395932.2 |  |
| *Nasonia vitripennis* | XP_001604969.2 | protein embryonic gonad | Nv XP_001604969.2 |  |
| *Pediculus humanus corporis* | XP_002430263.1 | conserved hypothetical protein | Phc XP_002430263.1 |  |
| *Pediculus humanus corporis* | XP_002430262.1 | conserved hypothetical protein | Phc XP_002430262.1 |  |
| *Pediculus humanus corporis* | XP_002430260.1 | knirps related protein, putative | Phc XP_002430260.1 |  |
| *Acyrtosiphon pisum* | XP_003243120.1 | knirps-related protein-like | Ap XP_003243120.1 |  |
| *Acyrtosiphon pisum* | XP_003248994.1 | protein embryonic gonad-like | Ap XP_003248994.1 |  |
| *Acyrtosiphon pisum* | XP_003244332.1 | protein doublesex* | Ap XP_003244332.1 |  |
| *Drosophila melanogaster* | NP_001259345.1 | ocelliless, isoform G | Dm Otd |  |
| *Tribolium castaneum* | NP_001034513.1 | orthodenticle-1 | Tc Otd-1 |  |
| *Tribolium castaneum* | KYB28866.1 | orthodenticle-2 | Tc Otd-2 |  |
| *Apis mellifera* | AER27696.1 | orthodenticle 1 | Am Otd-1 |  |
| *Apis mellifera* | AER27697.1 | orthodenticle 2 | Am Otd-2 |  |

## Acknowledgements

We thank Will Tank for technical assistance and helpful discussions. This work was supported by USDA-NIFA grant 2015-70016-23028, 2020-70029-33199 and HSI 1300394 in addition to an Institutional Development Award (IDeA) from the National Institute of General Medical Sciences of the National Institutes of Health under grant number P20GM103418.

## Author Contributions

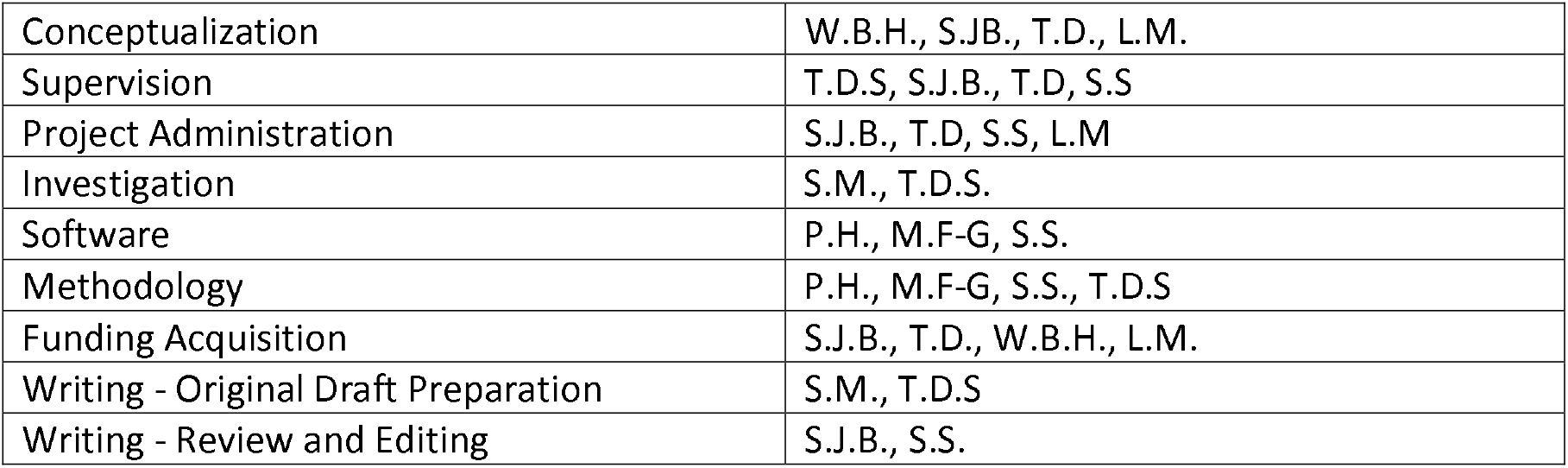

